# Oxaliplatin Induces Spontaneous Firing at Sensory Endings Across Touch and Proprioceptive Afferents

**DOI:** 10.64898/2026.07.03.736383

**Authors:** Paul Nardelli, J’Ana Reed, Jake Adam Vincent, Giulia A. Vitali, Kathy Clara Bui, Stephen Nicholas Housley, Timothy Charles Cope

## Abstract

Spontaneous activity in primary sensory neurons has been implicated in neuropathic symptoms, yet its earliest origins and immediate functional consequences remain incompletely understood. This gap is especially consequential in chemotherapy-induced peripheral neuropathy (CIPN), where sensory toxicities commonly limit effective cancer treatment. Using in vivo recordings in rats, we show that a single dose of oxaliplatin (OX) induces spontaneous firing within 24 h across touch and proprioceptive low-threshold mechanoreceptor (LTMR) afferents. Spontaneous firing consistently originated distally in peripheral axons and was accompanied by enhanced responses to mechanical stimulation, identifying LTMR sensory endings as the earliest source of spontaneous firing and a common site for spontaneous and stimulus-evoked hyperexcitability. OX also induced early structural abnormalities at sensory endings; however, SF+ LTMRs retained mechanosensory response profiles, indicating that spontaneous firing can emerge within otherwise functional sensory endings. Although coincident spontaneous and stimulus-evoked activity distorted encoding in individual LTMRs, these effects had little impact on population LTMR responses or motor behavior relying on mechanosensory feedback. Together, these findings identify sensory endings as an early target of OX neurotoxicity and demonstrate that spontaneous firing spanning multiple tactile and proprioceptive LTMR submodalities can coexist with largely preserved sensory function, indicating that even broad engagement across mechanosensory pathways is insufficient to disrupt all LTMR-dependent functions. These observations indicate that abnormal afferent activity initiated at sensory endings may be sufficient to engage sensory pathways underlying some paresthetic symptoms while leaving others largely unaffected, whereas progression to chronic neuropathic symptoms may require subsequent recruitment of the dorsal root ganglion.

**Highlights:**

- Oxaliplatin rapidly induces spontaneous firing across touch and proprioceptors.
- The earliest spontaneous firing localizes to sensory endings, not the DRG.
- Sensory endings are a shared site of spontaneous and evoked hyperexcitability.
- Spontaneous firing engages some sensory pathways while sparing others.

## Introduction

Primary somatosensory afferents normally generate action potentials only in response to specific somesthetic stimuli. Under pathological conditions, however, they can develop aberrant spontaneous firing, which has been implicated in a range of painful and non-painful sensations collectively associated with neuropathic disorders (Campero et al., 1998; Choi et al., 2024; Costigan et al., 2009; Devor, 2009; Finnerup et al., 2021; North et al., 2019; Raja et al., 2020; Smith, 2023). In chemotherapy-induced peripheral neuropathy (CIPN), spontaneous firing is thought to contribute to acute sensory symptoms, including dysesthesias and paresthesias (Cavaletti and Marmiroli, 2020; Flatters et al., 2017; North et al., 2018; Park et al., 2013; Wolf et al., 2012), which affect up to 95% of patients shortly after treatment (Argyriou et al., 2019; Pachman et al., 2015; Starobova and Vetter, 2017). These symptoms are often dose-limiting (Loprinzi et al., 2020) and predict the subsequent development and persistence of CIPN (Argyriou et al., 2013; Pachman et al., 2015). Defining the origins and functional consequences of spontaneous firing is therefore important for understanding disease progression and developing effective interventions.

Aberrant spontaneous firing may contribute to neuropathic symptoms through two non-exclusive mechanisms (Devor, 2009). First, sustained spontaneous activity may induce progressive neuroplastic changes that sensitize peripheral and central circuits, ultimately altering somatosensory processing (e.g., allodynia) (Campbell and Meyer, 2006; Devor, 2009; Latremoliere and Woolf, 2009; Raja et al., 2020). Alternatively, spontaneous firing may exert a more immediate effect by directly driving central sensory pathways through abnormal patterns of afferent activity (Chen et al., 2023; Devor, 2009; Zheng et al., 2022). In this direct-drive model, spontaneous firing can distort sensory signaling in real time by engaging sensory pathways that generate tingling, prickling, and numbness, without requiring the progressive neuroplastic changes associated with established neuropathy (Cavaletti and Marmiroli, 2020; Cavaletti et al., 2001; Gebremedhn et al., 2018; Nordin et al., 1984; Wolf et al., 2012). Support for this model comes from the close temporal relationship between spontaneous activity and symptom occurrence, together with the rapid relief of symptoms following local blockade of peripheral activity (Devor, 2009; Gracely et al., 1992; Pitcher and Henry, 2008; Smith, 2023). Despite substantial interest in spontaneous firing as a driver of neuropathic symptoms, its earliest peripheral origins and immediate functional consequences remain incompletely understood.

According to labeled-line theory (Finnerup et al., 2021), the perceptual consequences of spontaneous firing should depend on which sensory neuron subtypes generate aberrant activity. Spontaneous firing in large-diameter, fast-conducting afferents has long been associated with paresthesias in clinical studies (Ochoa and Torebjork, 1980; Xiao and Bennett, 2008), yet the relative involvement of functionally distinct low-threshold mechanoreceptor (LTMR) subtypes has not been systematically examined in vivo across cutaneous and proprioceptive populations (Campbell and Meyer, 2006; Choi et al., 2024; Saleque et al., 2024; Yamada et al., 2025). In addition, the trigger zone of chemotherapy-induced spontaneous firing remains unresolved (North et al., 2018). Spontaneous activity may arise at sensory endings, ectopically along peripheral axons, or within the dorsal root ganglion (DRG) (Amir et al., 2005; Han et al., 2023; Li et al., 2021; Nascimento et al., 2022; Zheng et al., 2022). Distinguishing among these possibilities is essential for refining mechanistic models of CIPN and identifying therapeutic targets. Clinical approaches such as microneurography cannot determine the site of SF initiation because recordings from individual neurons are limited to a single point along the axon.

Here we examined how oxaliplatin (OX), a standard-of-care chemotherapeutic associated with acute neurotoxicity (Ling et al., 2007; Saif and Reardon, 2005; Zajaczkowska et al., 2019), alters signaling in LTMR afferents. Using in vivo electrophysiological recordings in adult rats, we show that OX-induced spontaneous firing emerges rapidly, originates exclusively in the distal periphery, and occurs across all LTMR subtypes mediating touch and proprioception. We further demonstrate that spontaneous firing is accompanied by enhanced stimulus-evoked responsiveness, consistent with shared mechanisms underlying spontaneous and evoked hyperexcitability. Although spontaneous firing distorted stimulus encoding by individual LTMRs, its effects on population-level sensory signaling and proprioceptive behavior were surprisingly limited. These findings identify sensory endings as the earliest detected source of OX-induced spontaneous firing in LTMRs and reveal a dissociation between aberrant activity in individual afferents and impairment of sensory function during the acute phase of CIPN.

## Materials and Methods

### Animals

All procedures were approved by the Institutional Animal Care and Use Committees of Wright State University and the Georgia Institute of Technology. Experiments were performed on young adult female Wistar rats (250–300 g; Charles River Laboratories, Wilmington, MA, USA). Animals were housed under controlled temperature and light conditions with food and water available *ad libitum*. At the completion of terminal experiments, rats were euthanized by isoflurane overdose (5%) followed by exsanguination.

Rats were assigned to one of two experimental groups. Untreated control rats (n = 78) received no intervention. OX-treated rats (n = 50) received a single intraperitoneal (i.p.) injection of oxaliplatin (30 mg·kg⁻¹ in 5% dextrose) and were studied 24 h later. Preliminary dose-response experiments demonstrated that 30 mg·kg⁻¹ provided the most reliable detection of spontaneous firing (Supplementary Fig. S1A). Because this single-injection paradigm differs from those commonly used in preclinical OX studies (Calls et al., 2020; White et al., 2023; Xiao et al., 2012), DRG platinum accumulation was quantified as a surrogate measure of tissue OX exposure and compared with published preclinical and clinical measurements. An ex vivo calibration curve was also generated to relate commonly reported in vitro OX concentrations to DRG platinum accumulation (Supplementary Fig. S1B,C). Together, these analyses demonstrated that the selected dose provided a robust and clinically relevant framework for investigating the earliest peripheral mechanisms of OX-induced sensory dysfunction.

### In vivo electrophysiology

#### Surgical preparation

Experimental procedures have been described previously (Housley et al., 2022). Rats were anesthetized with isoflurane (1.5–2.5% in 100% O₂) throughout surgical preparation and data collection, which typically lasted approximately 8 h and concluded with euthanasia by isoflurane overdose followed by exsanguination. Core temperature (36–38°C), end-tidal CO₂ (3–5%), respiratory rate (40–60 breaths/min), heart rate (300–450 beats/min), and oxygen saturation (>90%) were monitored continuously and maintained by adjustment of anesthesia, radiant heat, and hourly subcutaneous saline administration.

Surgical procedures exposed selected skeletal muscles and peripheral nerves below the knee in the left hindlimb together with the ipsilateral L5 and L6 dorsal and ventral roots. Exposed tissues were covered with mineral oil throughout the experiment to prevent dehydration. Rats were secured in a stereotaxic frame using clamps attached to the snout, lumbar vertebrae, and distal femur of the left hindlimb to maintain a prone posture and provide stable conditions for electrophysiological recording and mechanical or electrical stimulation.

Skin and muscle LTMRs were studied in separate animals. For muscle recordings, the ankle was fixed, the Achilles tendon was severed at its insertion, and the triceps surae tendon was attached to the lever arm of a servomotor (Aurora Scientific Model 305C-LR) operated in length-servo mode to deliver computer-controlled muscle stretches (Housley et al., 2022). Corresponding peripheral nerves were dissected in continuity, separated from surrounding tissues, and positioned on bipolar stimulating electrodes within 5 mm of their muscle entry.

For skin recordings, the dorsum of the foot was secured to a platform, exposing the glabrous plantar surface to a rigid 1-mm probe driven by a servomotor operated in force-servo mode to deliver computer-controlled pressure stimuli (Housley et al., 2022). The posterior tibial nerve was isolated in continuity at the ankle and positioned on a bipolar stimulating electrode approximately 1.5 cm proximal to the center of the footpad.

#### Electrophysiological recording and LTMR classification

Action potentials were recorded intracellularly (Axoclamp 2B, Axon Instruments) from individual afferent axons using glass micropipettes (tip diameter ∼1 μm; 15–25 MΩ) filled with 3 M potassium acetate. Electrodes were advanced into dorsal roots in 1-μm steps using a microdrive (Transvertex). Primary sensory afferents were identified by orthodromic spikes evoked by electrical stimulation of the corresponding peripheral nerve (40 μs pulses, 1 pulse/s) and classified as Aβ low-threshold mechanoreceptors (LTMRs) based on conduction delays <3.5 ms.

Each afferent was studied as long as stable spike discrimination was maintained. LTMRs were classified as SF+ when spontaneous firing occurred in the absence of overt stimulation and SF− when it was absent. Controlled mechanical stimuli were then applied to classify functional LTMR subtypes. Classification was based on stimulus-evoked responses obtained between episodes of spontaneous firing or during periods of sparse, irregular firing (<10 spikes/s).

#### Muscle LTMRs

Group Ia and II muscle spindle afferents and group Ib tendon organ afferents were identified from their characteristic responses to electrically evoked muscle twitches and controlled muscle stretches (Vincent et al., 2017). Group Ia and II spindle afferents were distinguished from group Ib afferents by the pause in firing during muscle twitches evoked by peripheral nerve stimulation. Group Ia afferents were further distinguished from group II afferents by their high-fidelity responses to 100-Hz muscle vibration.

#### Cutaneous LTMRs

Receptive fields on the glabrous footpad were mapped using von Frey filaments. Responses to rectangular and sinusoidal pressure stimuli delivered in force-servo mode were used to classify afferents as rapidly adapting (RA) or slowly adapting (SA) (Handler and Ginty, 2021; Housley et al., 2022). RA afferents responded only at the onset and, in some cases, the offset of skin indentation. These criteria identified several RA afferents in which ongoing spontaneous firing could otherwise have resulted in classification as SA. Because subtype assignment remained uncertain, three cutaneous LTMRs were excluded from subtype analyses. Intermittent pauses in spontaneous firing allowed unequivocal classification of all remaining units. No attempt was made to distinguish subclasses of RA or SA afferents.

### Spike-triggered averaging

Spike-triggered averaging was used to localize the site of spontaneous firing initiation in intact LTMRs *in vivo*. Action potentials recorded from single dorsal root axons served as trigger events for temporal averaging of simultaneously recorded peripheral neurograms obtained with a custom-built extracellular amplifier. Averaging more than 50 triggered segments isolated time-locked peripheral events, allowing estimation of the site of spike initiation from the relative timing of trigger spikes and trigger-averaged responses (Fig. 4). Spike-triggered averages were unavailable for some LTMRs because recordings yielded insufficient trigger spikes or extracellular signals were degraded by saline accumulation around recording electrodes.

Counterfactual simulations predicted conduction delays expected if spontaneous firing originated at different locations along the LTMR pathway, including the peripheral recording site, the midpoint between recording sites, the DRG, and central recording sites. Forward models incorporated conduction velocities reported for skin and muscle LTMRs (Harper and Lawson, 1985; Leem et al., 1993; Lynn and Carpenter, 1982) together with conduction distances measured from experimental animals (Supplementary Fig. 2). The complete distribution from 1,000 simulations is reported. All code, priors, and raw data are available in the accompanying GitHub repository: https://github.com/housleylab/acute_oxaliplatin_encoding.

Electrical activity recorded from single axons and peripheral nerves, together with mechanical signals describing applied muscle stretch or skin pressure, was digitized (Cambridge Electronic Design), streamed continuously to a computer, and analyzed offline using Spike2 software.

### Immunohistochemistry and imaging

Triceps surae muscles (medial gastrocnemius, lateral gastrocnemius, and soleus) and glabrous skin from the left hindpaw were collected immediately after completion of terminal experiments. Tissues were fixed overnight in 4% paraformaldehyde at 4°C, cryoprotected in 30% sucrose, embedded in Tissue-Tek® Optimal Cutting Temperature (O.C.T.) compound (Sakura Finetek), and sectioned longitudinally at 50 μm using a Leica cryostat (Leica Microsystems, Buffalo Grove, IL). Sections were mounted on ColorFrost™ Plus glass slides (Epredia) and surrounded with a hydrophobic barrier.

Sections were washed three times (10 min each) in 0.01 M phosphate-buffered saline containing 0.3% Triton X-100 (PBS-T), blocked for 1 h in 10% normal goat serum diluted in PBS-T, incubated for 48 h at room temperature with rabbit anti-PGP9.5 antibody (1:500; Abcam, RRID: AB_10891773), washed, and incubated for 5 days at 4°C with Cy3-conjugated donkey anti-rabbit IgG (1:100; Jackson ImmunoResearch). Sections were washed again, mounted with VECTASHIELD® Antifade Mounting Medium containing DAPI (Vector Laboratories), coverslipped, sealed, and either imaged immediately or stored at 4°C.

Confocal imaging was performed using a Zeiss LSM 900 laser-scanning microscope equipped with a 63×/1.4 NA Plan-Apochromat oil-immersion objective and a 561-nm excitation laser. Z-stacks were acquired at 0.5-μm intervals and converted to average-intensity projections using Fiji (NIH). Figures were assembled using Adobe Illustrator or CorelDRAW.

For muscle spindle endings, the number of annulospiral turns, annulospiral width, and inter-spiral distance were quantified (Fig. 5). Inter-spiral distances from individual endings were averaged to generate a single measure of mean inter-spiral distance. Spindle endings containing fewer than five visible spirals were excluded to minimize variability arising from tissue sectioning.

For cutaneous sensory endings, epidermal innervation depth was quantified as the distance (μm) from the epidermal surface to the distalmost PGP9.5-positive terminal identified in maximum-intensity projection images. Five measurements were obtained from each animal and averaged to generate a single value for statistical analysis (n = 5 animals per group).

### Behavioral Analyses

Sensorimotor performance was assessed using the ladder-rung walking task, a validated measure of proprioceptive-guided locomotion (Metz and Whishaw, 2009), following previously described procedures (Housley et al., 2020b; Housley et al., 2021). Five rats were acclimated to the task for 3–4 days before baseline testing. Performance was assessed immediately before a single i.p. injection of OX and again 24 h later, immediately before terminal electrophysiological experiments, enabling within-subject comparisons of pre- and post-treatment performance.

Rats traversed a 10° decline with unevenly spaced ladder rungs while locomotion was recorded using a GoPro Hero12 Black camera (30 frames/s). More than 100 left hindlimb steps (102–128) were analyzed from five walking trials for each animal. Analyses focused on the left hindlimb because afferent encoding was subsequently examined electrophysiologically in that limb.

Healthy rats normally place the ipsilateral hindpaw on the same rung previously contacted by the ipsilateral forepaw, referred to here as the hindlimb replacement strategy (Housley et al., 2021; Metz and Whishaw, 2009; Vincent et al., 2016). Hindlimb performance was quantified from sagittal-plane video recordings using two complementary measures: incorrect rung replacement, defined as placement of the hindpaw on a rung other than the intended target rung, and failure to secure rung contact, defined as failure to establish or maintain secure contact with any rung, including missed contacts, atypical foot placement, and slips. Error rates were calculated from all steps for each rat.

Video recordings were scored independently by two investigators blinded to treatment group, yielding substantial inter-rater agreement (Cohen’s κ = 0.71). Error rates were averaged across trials to obtain a single value for each animal. In total, there were 43 agreements out of 50 observations.

## Statistical analyses

Statistical analyses of neuronal encoding have been described previously (Horstman et al., 2019; Housley et al., 2020a). Bayesian parameter estimation was used to derive the joint posterior distribution of all model parameters simultaneously. Highest posterior density intervals (95% HDIs) were used to compare posterior probability distributions between contrasts of interest (Horstman et al., 2019; Housley et al., 2020a). Models were implemented using the rstanarm package (v2.18.1) (Gabry and Goodrich, 2018) in R (v3.5.0) (Team, 2018). Model performance was evaluated using leave-one-out cross-validation with Pareto-smoothed importance sampling (PSIS) (Vehtari et al., 2017), as described previously (Housley et al., 2020a). Summary statistics are reported as mean ± SE unless otherwise indicated.

## Results

### I. Acute OX-induced spontaneous firing in LTMRs

#### Spontaneous firing patterns and incidence

Figure 1 illustrates representative examples of spontaneous firing observed within 24 h of a single i.p. injection of OX. SF occurred in LTMRs innervating both skeletal muscle and glabrous skin and exhibited two distinct patterns. The predominant pattern consisted of irregular firing with highly variable inter-spike intervals (Fig. 1B). A less common pattern comprised bursts of rapid firing lasting 1–15 s with recognizable, although variable, periodicity (Fig. 1C). Similar firing patterns have been reported previously in Aβ afferents following exposure to OX, paclitaxel, and vincristine (Choi et al., 2024; Xiao and Bennett, 2008; Xiao et al., 2012).

**Figure 1.**
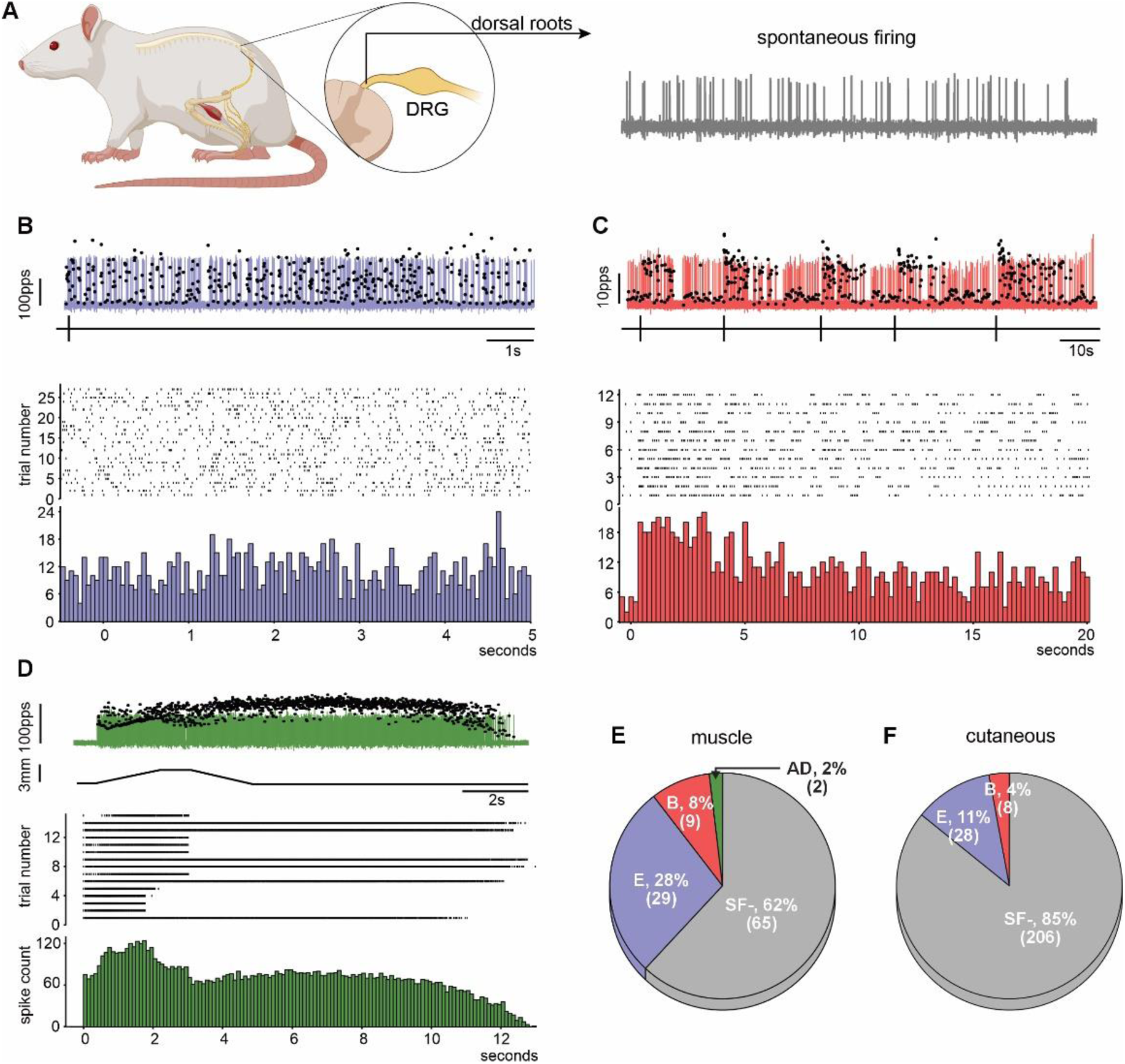
Patterns and incidence of spontaneous firing induced by acute OX in LTMRs. **(A)** Illustration of the dorsal root recording site proximal to the dorsal root ganglion together with a representative recording showing spontaneous firing. **(B–D)** Representative patterns of spontaneous firing and afterdischarge in LTMRs. In each panel, the upper trace shows action potentials superimposed with black dots indicating instantaneous firing rate; the middle trace shows a raster plot; and the lower trace shows the corresponding spike histogram. **(B)** Erratic spontaneous firing (blue) in a muscle LTMR, characterized by irregular firing and a diffuse spike histogram. **(C)** Bursting spontaneous firing (red) in a skin LTMR, characterized by episodes of relatively high-frequency firing. Increased spike density at burst onset is evident in the histogram and aligned with the vertical black lines beneath the upper trace. **(D)** Afterdischarge (green), defined as firing that persists after termination of a muscle stretch stimulus in a muscle LTMR. **(E, F)** Pie charts showing the proportions and numbers of muscle (E) and skin (F) LTMRs exhibiting erratic spontaneous firing (blue), bursting spontaneous firing (red), and afterdischarge (green), or no spontaneous firing (SF-, gray). Data were collected from 21 rats for muscle LTMRs and 29 rats for skin LTMRs.

A rare form of abnormal activity, designated afterdischarge, was also classified as SF+. In these LTMRs, mechanically evoked firing persisted after stimulus termination and, in some trials, exceeded firing rates observed during stimulation (Fig. 1D). Afterdischarge was never observed in untreated rats, occurred only rarely in muscle LTMRs following OX treatment (2%), and was not detected in skin LTMRs.

LTMRs lacking spontaneous activity during the recording period were classified as SF− and constituted the majority of units recorded from both muscle and skin (Figs. 1E,F). Because SF was often intermittent (see below), the measured incidence likely underestimated its occurrence over longer recording periods. When grouped by tissue, the proportion of SF+ LTMRs was approximately threefold greater in muscle than in skin. Nevertheless, SF incidence in both tissues was markedly elevated relative to untreated rats, in which spontaneous activity was rare (<1%; 2/142), based on a compiled database from multiple studies (Housley et al., 2022; Vincent et al., 2017). These observations are consistent with previous reports of acute CIPN models (Choi et al., 2024; Han et al., 2023; Xiao et al., 2012).

Because SF was intermittent, incidence alone did not fully describe its magnitude. Raster plots (Figs. 2A,B) illustrate the temporal organization of SF in individual muscle and skin LTMRs, revealing substantial variability among afferents. Expanded views highlight the episodic nature of SF. Figures 2C,D quantify the proportion of recording time occupied by erratic firing, bursting, afterdischarge, or silence.

**Figure 2.**
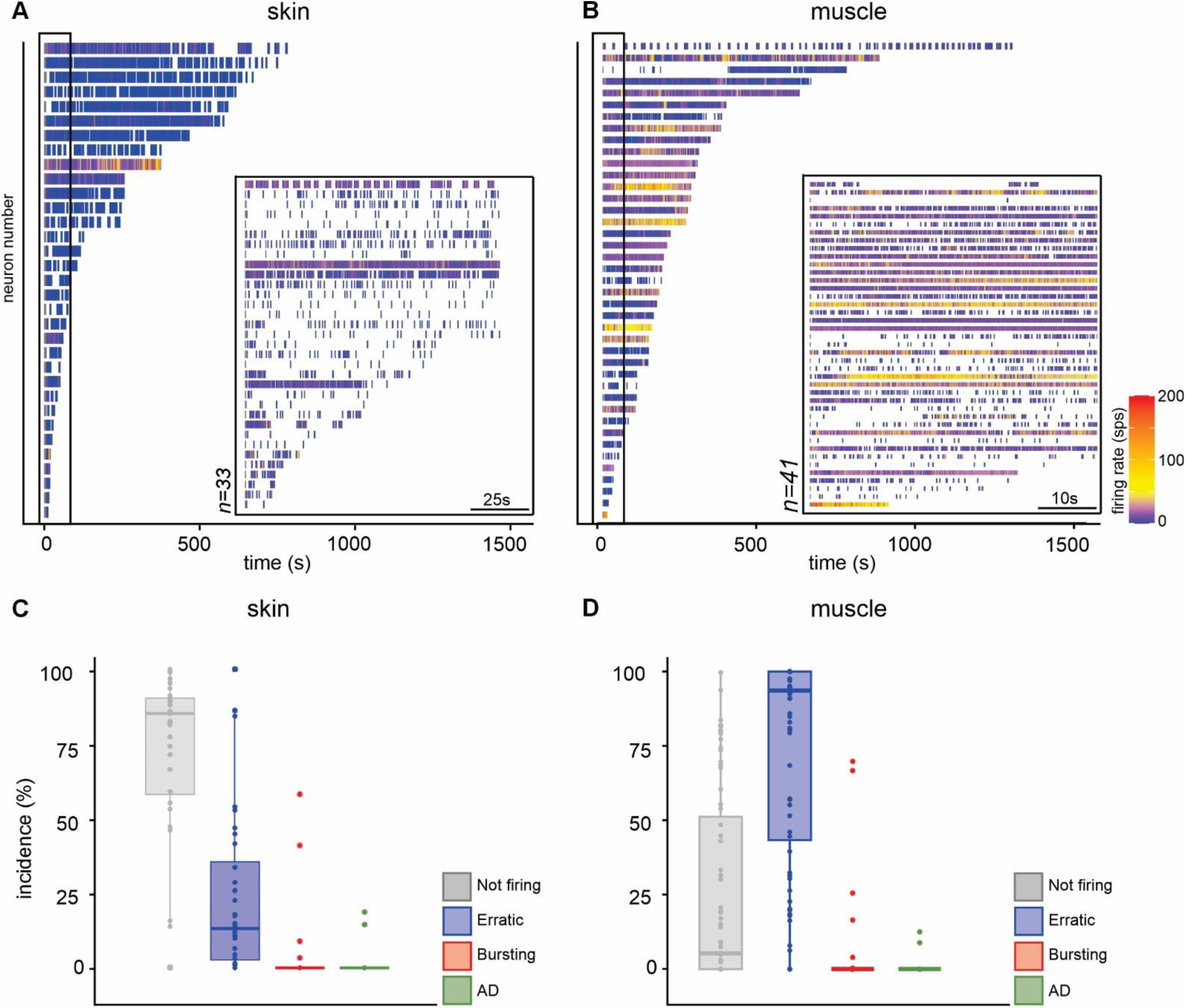
Duration of spontaneous firing in LTMRs. **(A, B)** Raster plots showing bouts of spontaneous firing recorded from individual LTMRs in skin **(A)** and muscle **(B)**. LTMRs are ranked from top to bottom by the total duration of spontaneous firing; recording duration varied among units because of differences in axonal recording stability. Insets show time-expanded regions corresponding to the boxed areas; note that the inset time scales differ. **(C, D)** Duration of spontaneous firing expressed as the percentage of recording time occupied by spontaneous firing. Box-and-whisker plots show the distribution of recording time spent in erratic, bursting, afterdischarge, or silent states by SF+ LTMRs in skin **(C)** and muscle **(D).** Boxes indicate quartiles and medians; whiskers and points indicate the range and outliers.

### Spontaneous firing across LTMR subtypes

We next asked whether the higher incidence of spontaneous firing in muscle than skin reflected differences among LTMR subtypes. Across repeated stimulus trials, the firing patterns used to classify LTMR subtypes (see Methods) remained stable when reassessed during periods in which SF was absent. SF did not alter the physiological criteria used to identify LTMR subtypes.

Figures 3A and 3B show that approximately one-third of group II muscle spindle afferents, group Ib tendon organ afferents, and slowly adapting (SA) skin afferents exhibited SF+. The overall difference between muscle and skin LTMRs was largely attributable to two subtypes. Nearly twice as many group Ia spindle afferents exhibited SF+ as the other muscle LTMR subtypes, whereas only 10% of rapidly adapting (RA) skin afferents exhibited SF+ activity.

**Figure 3.**
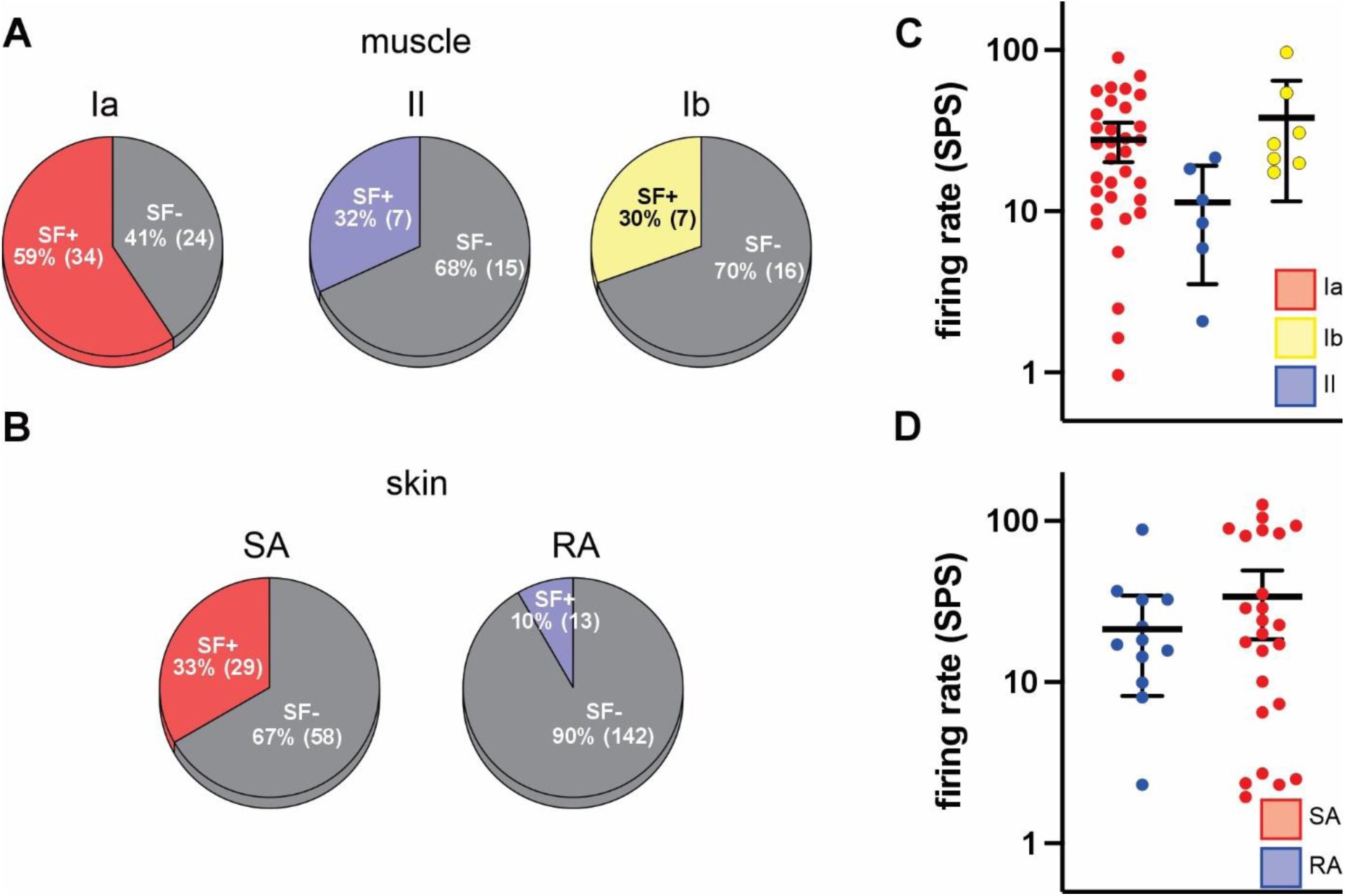
Incidence and firing rates of spontaneous activity across LTMR subtypes. **(A,B)** Pie charts showing the proportions and numbers of muscle **(A)** and skin **(B)** LTMR subtypes exhibiting SF+. **(C, D)** Box-and-whisker plots comparing SF+ firing rates across muscle (C) and skin (D) LTMR subtypes. Instantaneous firing rates were averaged over 30-s intervals corresponding to the shortest spontaneous firing episode observed in the dataset. Plots show medians, interquartile ranges, and outliers for individual LTMRs within each subtype.

The low incidence of SF+ among RA afferents raised the possibility that OX selectively suppressed RA responsiveness, as reported previously in ex vivo skin–nerve preparations (Saleque et al., 2024) and after repeated vincristine treatment (Yamada et al., 2025). This explanation appears unlikely because the relative proportions of SA and RA afferents identified by their stimulus-response properties were similar in untreated (66% SA) and OX-treated (57% SA) animals.

Figures 3C and 3D compare SF firing rates across LTMR subtypes. Instantaneous firing rates were averaged over 30-s intervals, corresponding to the shortest SF+ episode observed in the dataset. Median firing rates varied among subtypes but generally exceeded 10 spikes/s.

### II. Spontaneous firing originates in distal LTMR axons early after OX treatment

#### Acute spontaneous firing originates distal to the peripheral recording site

The cellular origin of chemotherapy-induced SF remains unresolved (North et al., 2018). To identify the earliest source of acute SF, we used spike-triggered averaging (Methods; Fig. 4A–D) to compare action potential arrival times at central and peripheral recording sites in fully intact LTMRs *in vivo*. This approach avoided experimental manipulations, such as axonal transection, that can themselves induce spontaneous firing (Michaelis et al., 2000).

Figures 4E and 4F illustrate the temporal relationship between spikes recorded in dorsal roots and their corresponding mechanically evoked or spontaneous spikes detected in the peripheral nerve. Across all LTMRs examined (Figs. 4G,H), spontaneous spikes recorded in the peripheral nerve consistently preceded their arrival at the central recording site by delays matching axonal conduction between the two recording sites. These observations localize spontaneous firing to a site at or distal to the peripheral recording electrode across all LTMR subtypes.

To further constrain the site of spike initiation, we modeled the conduction delays expected for spikes arising from six anatomically plausible locations along the LTMR pathway (Fig. 4I) using published conduction velocities and measured axonal lengths (Supplementary Fig. 2). Only simulations assuming spike initiation distal to the peripheral recording site reproduced the experimentally observed timing. For example, spikes originating in the DRG would be expected to arrive at the peripheral recording electrode after detection at the central recording site, opposite to the observed pattern. These findings localize spontaneous firing during the first 24 h after OX treatment to the distal periphery rather than the DRG or more proximal axonal locations.

Although these experiments did not identify the precise site of spike initiation, additional observations were consistent with an origin at or near sensory endings. Lidocaine applied to the nerve distal to the recording electrode abolished spontaneous firing in five rats across multiple LTMR subtypes (three Ia, one II, and one SA). Similarly, spontaneous firing was eliminated within 2 min of applying lidocaine directly to the muscle surface over the receptive field of a group Ia afferent (Fig. 4J). Stretch-evoked firing persisted in other LTMRs recorded simultaneously from the same dorsal root (Fig. 4K), and electrically evoked muscle contractions remained intact (Fig. 4L), indicating that the local anesthetic block was spatially restricted.

**Figure 4.**
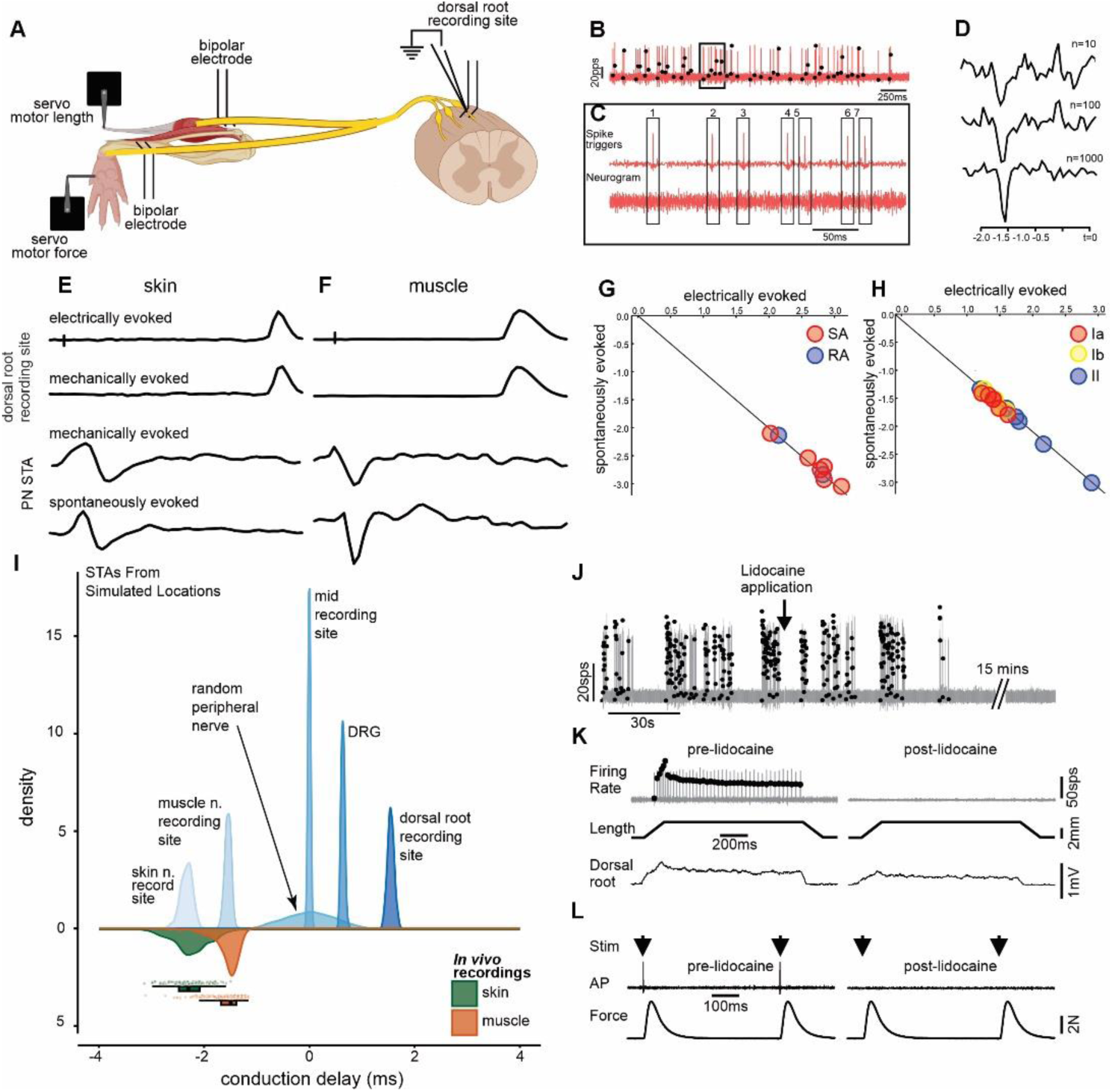
Distal origin of OX-induced spontaneous firing. **(A)** Experimental arrangement for recording single-axon spikes in dorsal roots and using those events to trigger time-locked averaging of activity recorded in peripheral nerves (B–D; see Methods). **(E,F)** Representative recordings from skin **(E)** and muscle **(F)** LTMRs aligned vertically in time. The upper two traces show activity recorded centrally in dorsal roots (DR) from individual axons, including spikes evoked by electrical stimulation of peripheral nerves or mechanical stimulation of receptive fields. The lower two traces show spike-triggered average (STA) events recorded at the peripheral nerve (PN) site preceding the central trigger spikes by delays matching those of peripherally evoked spikes. **(G,H)** Conduction times between peripheral and central recording sites for spontaneous and electrically evoked activity plotted for all LTMRs by subtype. The line of identity indicates equivalent conduction times for spontaneous and evoked spikes. Longer conduction times for skin LTMRs reflect the greater distance between peripheral and central recording sites (A; Supplemental Fig. 1). **(I)** Simulated conduction time distributions for spontaneous firing originating at different locations along the LTMR pathway, derived from measured conduction velocities and estimated axonal lengths (see Supplemental Figure 2). Times are referenced to zero, defined as the moment when spikes originating midway between recording sites would arrive simultaneously at both sites. Only simulations assuming spike initiation distal to the peripheral recording site reproduced the observed conduction delays. **(J-L)** Lidocaine applied to the receptive field of a muscle spindle Ia LTMR selectively abolished spontaneous firing **(J)** while preserving evoked activity in other LTMRs, as indicated by maintained dorsal root responses to muscle stretch **(K)** and intact motor axon conduction, confirmed by persistent muscle twitch contractions **(L)**.

### III. Early structural changes at peripheral sensory endings

To identify structural abnormalities associated with SF, we examined sensory endings from LTMR populations exhibiting the highest incidence of spontaneous activity. Muscle spindle endings of group Ia afferents displayed heterogeneous morphology following OX treatment. Within the same animal, some annulospiral endings appeared indistinguishable from controls, whereas others exhibited conspicuous thinning and disorganization (Fig. 5A–C). Two blinded observers showed substantial agreement when classifying spindle endings as normal or abnormal, following OX treatment (Cohen’s κ = 0.74). In this analysis, there were 46 agreements out of 50 total observations Quantitative analysis showed no difference in the number of annulospiral turns between untreated and OX-treated animals (15.0 ± 1.3 vs. 12.6 ± 1.0). In contrast, both annulospiral width and mean inter-spiral distance differed significantly between groups (Fig. 5D–F).

**Figure 5.**
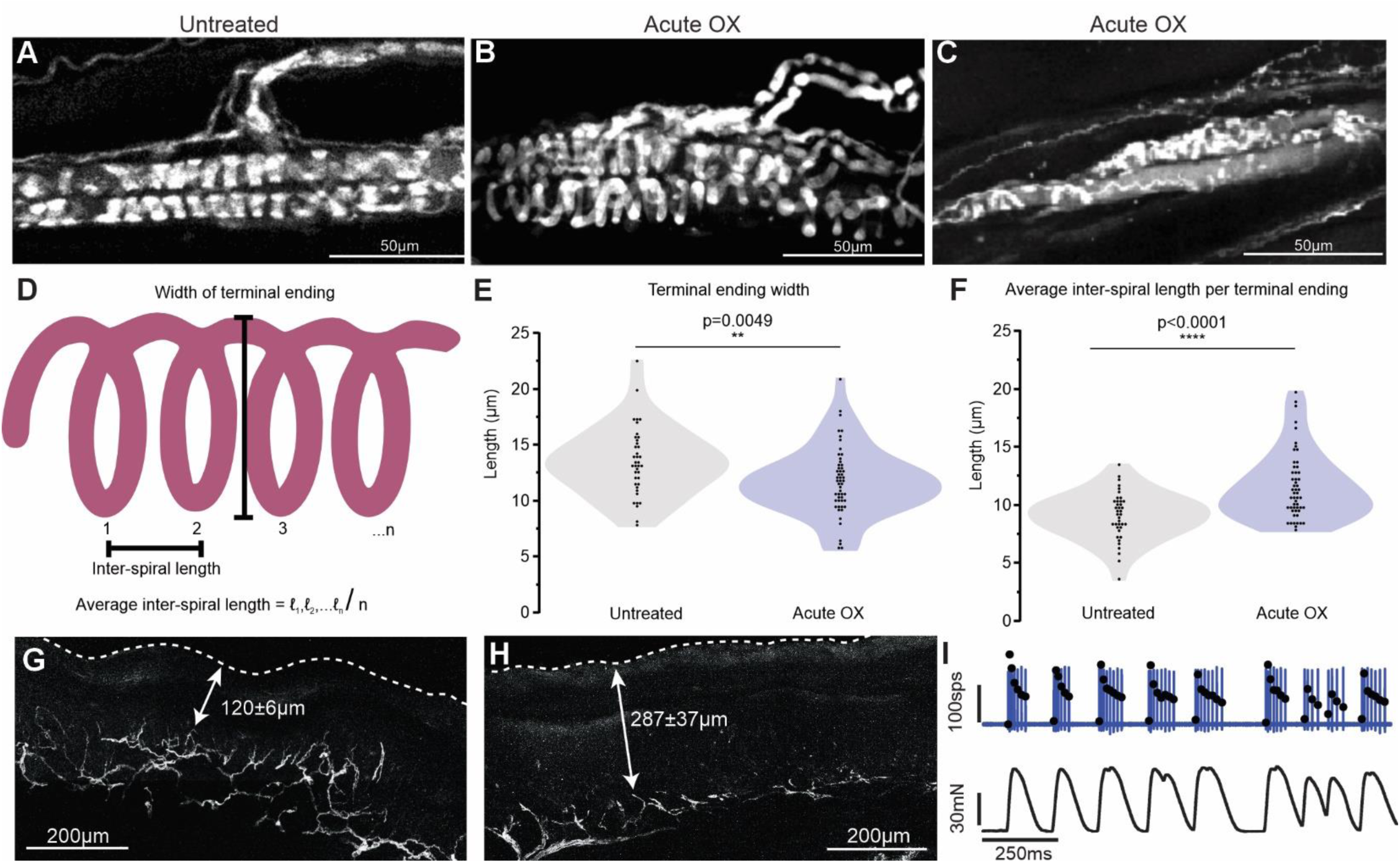
Early peripheral consequences of acute oxaliplatin (OX) exposure. Representative confocal images of PGP9.5 immunoreactivity showing sensory endings in medial gastrocnemius (MG) muscle spindles (A–C) and glabrous skin (G,H) from untreated and OX-treated rats. (A) Normal Ia afferent annulospiral ending in an untreated rat. (B,C) Representative Ia afferent endings from OX-treated rats exhibiting either a normal appearance (B) or thinning and disorganized annulospiral projections (C). (D) Schematic illustrating measurements used to quantify annulospiral morphology. Violin plots summarize annulospiral width (E) and inter-spiral distance (F) measured from 39 muscle spindles in 6 untreated rats and 53 muscle spindles in 3 OX-treated rats. Although considerable overlap was observed between groups, OX treatment significantly reduced annulospiral width and increased inter-spiral distance. (G,H) Representative images of glabrous skin showing reduced PGP9.5 immunoreactivity, thinning, and retraction of epidermal sensory nerve terminals after OX treatment. Quantification demonstrated a marked reduction in epidermal innervation depth in OX-treated rats compared with untreated rats (120 ± 6 µm vs. 287 ± 37 µm; n = 5 per group). (I) Representative recording illustrating spontaneous muscle fasciculations that directly activated a group Ib afferent.

Recordings from muscle LTMRs identified an additional peripheral source of abnormal activity. Simultaneous force recordings obtained from 18 muscle LTMRs detected spontaneous muscle fasciculations, a recognized consequence of OX treatment (Hill et al., 2010; Lehky et al., 2004). Figure 5G illustrates a representative example in which a fasciculation elicited firing in a group Ib afferent. Spike-triggered averaging of muscle force demonstrated that abnormal firing in five group Ib afferents and one group II afferent, initially classified as SF+, was attributable to fasciculations. Fasciculations also modulated spontaneous firing in five group Ia afferents, although spontaneous firing persisted independently. Weak muscle contractions (<1 mg), likely reflecting fibrillations, were not associated with LTMR firing.

Cutaneous sensory endings also exhibited structural abnormalities after OX treatment. Epidermal nerve terminals showed thinning, and retraction compared with untreated animals (Figs. 5H,I). Quantitative analysis confirmed a marked reduction in epidermal innervation depth (120 ± 6 μm vs. 287 ± 37 μm; n = 5). These observations are consistent with previous reports demonstrating structural changes within 24 h of OX treatment (Xiao et al., 2012).

Structural abnormalities emerged at both cutaneous and proprioceptive sensory endings within 24 h of OX administration. These findings identify sensory endings as an early target of OX neurotoxicity while supporting their role as a potential source of spontaneous firing (Caudle and Neubert, 2022).

### IV. Acute OX enhances mechanically evoked LTMR responses

We hypothesized that if spontaneous firing reflects increased excitability of LTMR sensory endings, it should be accompanied by enhanced responsiveness to mechanical stimulation, providing one potential explanation for the tactile hypersensitivity characteristic of acute OX-induced neuropathy.

Figure 6 shows representative examples of mechanosensory encoding together with the response parameters quantified for muscle and skin LTMRs. Overall firing profiles resembled those reported previously in untreated rats (Housley et al., 2022; Vincent et al., 2017), but consistent differences emerged following OX treatment. Hierarchical Bayesian modeling was used to determine whether spontaneous firing classification (SF+ or SF−) predicted changes in mechanically evoked responses.

**Figure 6.**
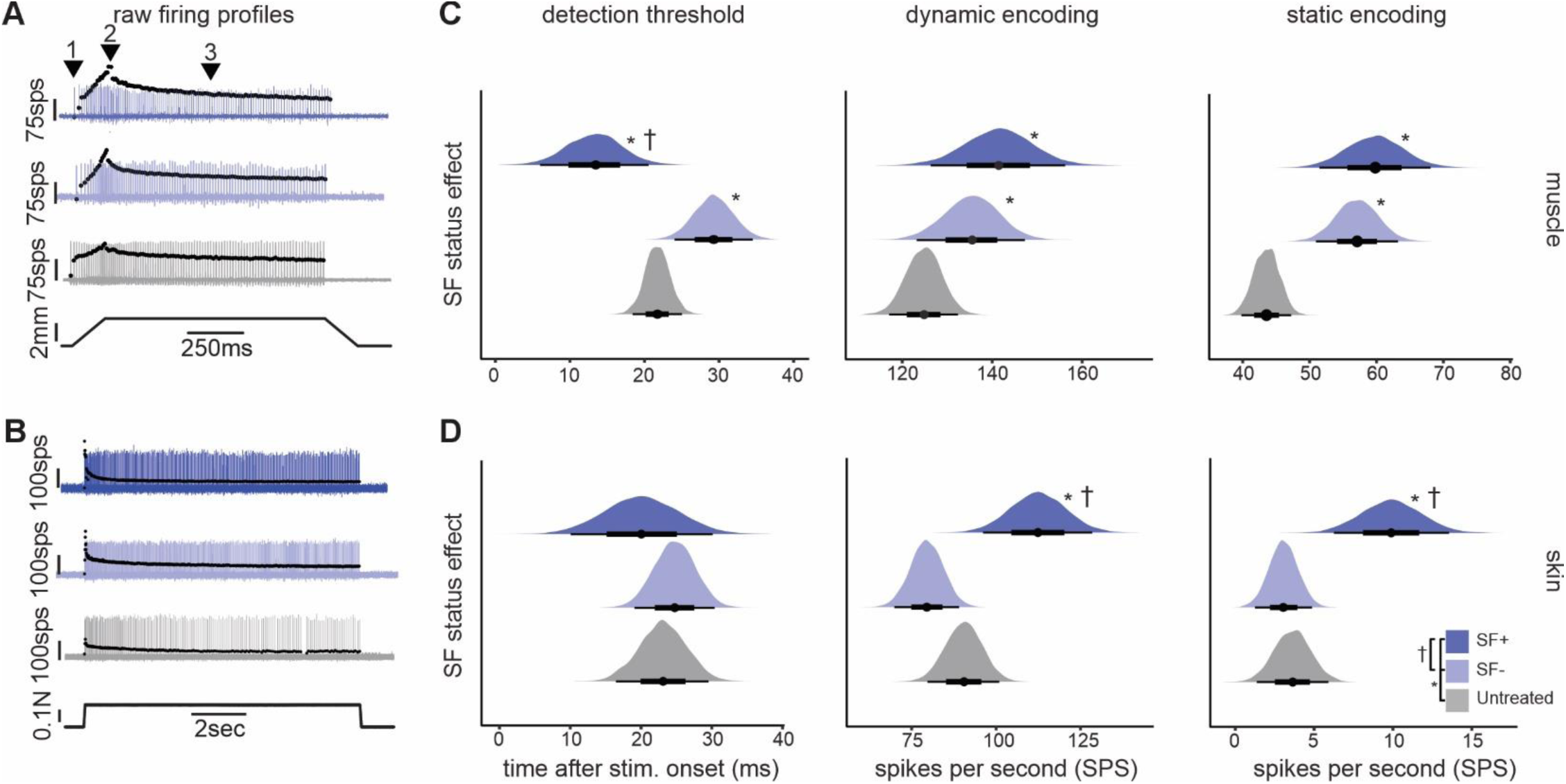
Enhanced mechanosensory responsiveness following OX exposure. **(A, B)** Representative firing responses to muscle stretch and skin pressure by LTMRs from control and OX-treated rats, classified as SF+ or SF-. Numbered arrows in (A) indicate firing parameters quantified for both muscle and skin LTMRs: (1) detection threshold, defined by the first spike after stimulus onset; (2) dynamic encoding, measured as the instantaneous firing rate at the peak of increasing stretch or pressure; and (3) static encoding, quantified as the instantaneous firing rate at the stimulus midpoint. **(C,D)** Hierarchical Bayesian modeling assessing the influence of SF⁺/SF⁻ classification on LTMR mechanosensory responses. Significant effects indicate altered mechanotransduction and/or spike-encoding properties following OX treatment.

Figures 6C and 6D show that OX significantly enhanced LTMR responsiveness. Relative to untreated LTMRs, SF+ LTMRs exhibited lower firing thresholds together with increased dynamic and static firing rates. These effects reached statistical significance in both muscle and skin LTMRs, except for firing threshold in skin afferents. SF+ LTMRs therefore exhibited both spontaneous firing and enhanced responsiveness to natural mechanical stimuli.

Because responses differed among SF+ and SF− populations, we next examined whether particular LTMR subtypes accounted for these effects. Subtype-specific analyses identified group Ia spindle afferents as the principal contributors to enhanced responsiveness in both SF+ and SF− populations, whereas group Ib afferents contributed minimally. Because γ-motoneuron activation increases group Ia sensitivity, we considered whether OX indirectly enhanced spindle responsiveness through the fusimotor system. Ventral root transection, which eliminates γ-motoneuron drive to muscle spindles (Matthews, 1972), did not alter group Ia mechanosensory encoding, indicating that the enhanced responsiveness was not attributable to fusimotor activation.

### V. Spontaneous firing disrupts individual but not population encoding

#### Encoding by individual LTMRs

Recordings from individual LTMRs demonstrated that spontaneous firing directly disrupted the representation of mechanical stimuli. Representative examples (Figs. 7A–C) compare stimulus encoding during episodes of spontaneous firing with responses recorded from the same afferents during periods when spontaneous activity was absent.

**Figure 7.**
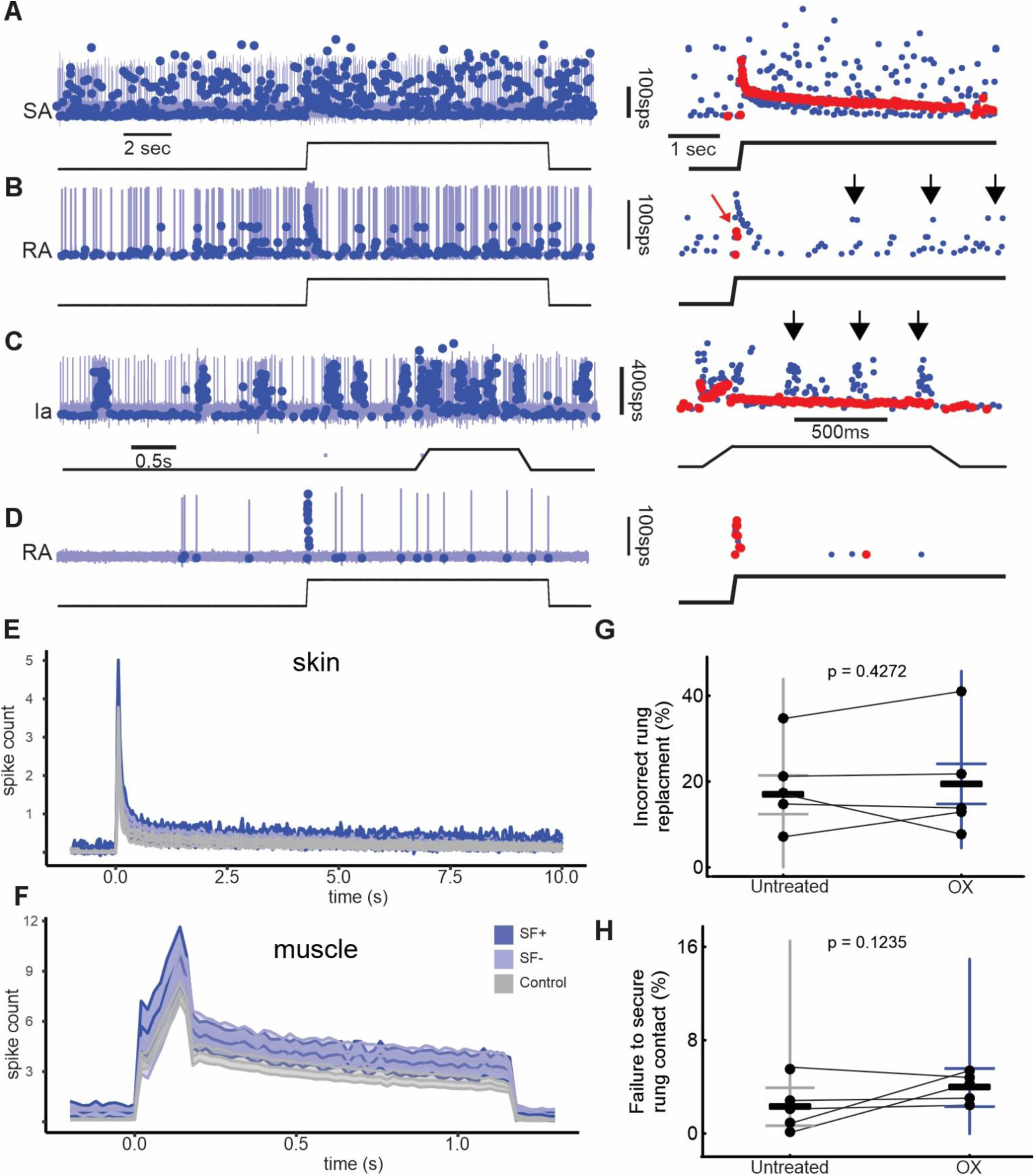
Spontaneous firing disrupts individual LTMR encoding while preserving population encoding and sensorimotor function. **(A–D)** Representative examples showing interactions between spontaneous firing and stimulus-evoked spike encoding in individual LTMRs. Blue vertical lines and blue dots indicate action potentials and corresponding instantaneous firing rates, respectively, occurring before and during applied muscle stretch or skin pressure (lower traces). Expanded views (right) compare instantaneous firing rates during spontaneous firing (blue dots) with stimulus-evoked responses recorded during separate trials in the absence of spontaneous firing (red dots), illustrating disruption of stimulus encoding by spontaneous activity. **(A)** Erratic spontaneous firing reduces the signal-to-noise ratio during skin-pressure encoding in a slowly adapting (SA) skin LTMR. **(B)** In a rapidly adapting (RA) skin LTMR, SF+ generates spurious signals of repeated stimulus onset (arrows) in the absence of changes in skin pressure. **(C)** In a muscle spindle Ia LTMR, burst firing obscures encoding of the dynamic phase of muscle stretch and generates inappropriate dynamic-like firing during the static hold phase of stretch (arrows). **(D)** Sparse, irregular SF+ has little impact on encoding of skin-displacement onset in an RA skin LTMR. **(E,F)** Population-level encoding was assessed by aligning responses from multiple LTMRs to identical mechanical stimuli and averaging across afferents. LTMRs from OX-treated rats were grouped as SF+ or SF-, with untreated controls shown for comparison. Population responses are shown for the LTMR subtypes with the highest incidence of spontaneous firing: slowly adapting skin LTMRs (E) and muscle spindle Ia LTMRs (F). Despite enhanced mechanosensory responsiveness in SF+ afferents (cf. Fig. 5), population-averaged encoding remained highly similar across SF+, SF−, and control groups, with root-mean-square deviations <5%. **(G,H)** Longitudinal assessment of sensorimotor performance before and 24 h after a single i.p. injection of OX. Error rates are expressed as the percentage of incorrect rung replacement (G) or failure to secure rung contact (H) during more than 100 steps recorded from each rat over five walking trials. Symbols connected by lines represent data from individual rats. Group mean performance and individual-animal performance remained within the range of baseline variability measured before treatment, indicating that acute OX-induced spontaneous firing did not impair this LTMR-dependent sensorimotor behavior. Vertical bars indicate the group range, and horizontal bars denote the mean ± confidence interval.

In a slowly adapting cutaneous LTMR (Fig. 7A), spontaneous firing reduced the signal-to-noise ratio, partially masking skin-displacement signals and degrading encoding fidelity. In a rapidly adapting cutaneous LTMR (Fig. 7B), spontaneous firing amplified responses to dynamic skin displacement and generated inappropriate firing during static displacement, producing spurious dynamic signals. In a group Ia spindle afferent (Fig. 7C), spontaneous firing obscured encoding of the dynamic phase of muscle stretch and generated dynamic-like firing during the subsequent static hold phase. These examples demonstrate that spontaneous firing can mask, distort, or generate false mechanosensory signals across multiple LTMR subtypes. In contrast, stimulus encoding remained largely preserved when spontaneous firing occurred infrequently and at low firing rates (Fig. 7D).

#### Encoding by LTMR populations

We next asked whether these abnormalities at the level of individual afferents altered population encoding. Figures 7E and 7F show ensemble responses constructed from representative muscle and skin LTMR subtypes. Analyses focused on group Ia muscle spindle afferents and slowly adapting cutaneous afferents because these subtypes exhibited the highest incidence of SF+ activity (Fig. 3A,B).

Despite the pronounced effects of spontaneous firing on individual LTMRs, population responses from OX-treated rats closely resembled those from untreated rats and differed little between SF+ and SF− groups. The principal effect of spontaneous firing was a modest shift toward slightly earlier and more variable firing. Consequently, the distortions observed in individual afferents (Figs. 7A–D) had little effect on population-level encoding of mechanical stimuli. The predominance of SF-afferents together with the intermittent and asynchronous expression of SF+ activity effectively smoothed the aggregate signal and preserved population representations of mechanical stimuli.

### VI. Preserved sensorimotor performance despite acute OX-induced spontaneous firing

To determine whether OX-induced spontaneous firing was associated with measurable sensorimotor deficits, we longitudinally assessed performance on a ladder-rung walking task that depends on tactile and proprioceptive feedback and is sensitive to LTMR dysfunction in chronic CIPN models (Han et al., 2023; Vincent et al., 2016). This assay was selected because it provides a direct assessment of LTMR-dependent sensorimotor function during a natural locomotor behavior [7].

Figures 7G and 7H summarize longitudinal changes in performance of the hindlimb replacement strategy before and 24 h after a single i.p. injection of OX. Across more than 100 steps per animal (102–128 steps over five walking trials), both incorrect rung replacement (Fig. 7G) and failure to secure rung contact (Fig. 7H) remained within the range of baseline variability for each animal, indicating stable sensorimotor performance following OX treatment.

Electrophysiological recordings obtained immediately after behavioral testing confirmed the presence of SF+ activity and enhanced mechanosensory responsiveness in the same animals. Preservation of the hindlimb replacement strategy therefore occurred despite acute abnormalities in LTMR signaling.

Acute OX-induced spontaneous firing and enhanced LTMR responsiveness were not accompanied by impairment of this LTMR-dependent behavior. This observation is consistent with the preserved population-level encoding described above and indicates that abnormal activity distributed across multiple tactile and proprioceptive LTMR subtypes is insufficient to disrupt all LTMR-dependent functions.

## Discussion

Our study identifies sensory endings as the earliest detected source of oxaliplatin-induced spontaneous firing in low-threshold mechanoreceptors and provides new insight into how this activity alters sensory signaling during the acute phase of chemotherapy-induced peripheral neuropathy (CIPN). We show that a single dose of OX induces spontaneous firing within hours across multiple LTMR subtypes mediating touch and proprioception. Using *in vivo* recordings from intact afferents, we localize the site of spike initiation to distal regions of the peripheral axon adjacent to sensory endings and demonstrate that spontaneous firing is accompanied by enhanced stimulus-evoked responsiveness. These observations identify sensory endings as a common site of both spontaneous and evoked hyperexcitability.

Despite marked distortion of spike patterns in individual LTMRs, OX-treated rats retained normal performance of the hindlimb replacement strategy during a sensorimotor task dependent on tactile and proprioceptive feedback. This dissociation between abnormal activity in individual afferents and preserved population-level encoding and sensorimotor performance reveals an important limitation of spontaneous firing during the acute phase of OX exposure. More broadly, our findings indicate that abnormal activity distributed across tactile and proprioceptive LTMR submodalities is insufficient to disrupt all LTMR-dependent pathways. Instead, the functional consequences of spontaneous firing appear to depend on how aberrant activity is integrated within specific central circuits, allowing some sensory pathways to be engaged while others remain largely unaffected.

### Sensory endings are the earliest identified source of spontaneous firing in LTMRs

Previous studies reported spontaneous firing in peripheral sensory axons following repeated exposure to chemotherapeutic agents, including OX, paclitaxel, and vincristine (Xiao et al., 2012; Xiao et al., 2011). Our findings extend this work by demonstrating that spontaneous firing emerges within hours of a single OX exposure. Using recordings from intact afferents in continuity with the DRG, we show that this early spontaneous firing originates consistently and exclusively within distal axons adjacent to sensory endings. These findings identify sensory endings as the earliest detected source of spontaneous firing and suggest that they are among the earliest targets of OX neurotoxicity.

Consistent with this localization, OX produced rapid structural abnormalities at sensory endings in both skin and muscle. In glabrous skin, we confirmed and extended previous reports of thinning, retraction, and loss of intraepidermal sensory terminals following OX exposure (Caudle and Neubert, 2022; Siau et al., 2006; Xiao et al., 2012), demonstrating that these abnormalities can emerge within 24 h. We further show that proprioceptive endings within muscle spindles undergo similarly rapid structural remodeling. Early terminal pathology therefore extends beyond cutaneous afferents and appears to represent a common response across multiple LTMR subtypes.

Despite these structural abnormalities, mechanosensory function remained largely preserved. All SF+ LTMRs retained characteristic response properties and could be classified according to established physiological criteria, indicating that mechanotransduction and spike encoding remained robust despite evidence of terminal remodeling. Spontaneous firing can therefore emerge within structurally disrupted yet functional sensory endings, indicating that terminal pathology reflects more than passive degeneration.

Instead, sensory endings remain capable of generating aberrant activity during the earliest stages of OX exposure. A similar dissociation between structural abnormalities and preserved sensory function has been reported in regenerated muscle spindle afferents (Barker et al., 1985)

### Shared terminal mechanisms may underlie spontaneous and evoked hyperexcitability

One of the most striking observations was the close association between spontaneous firing and enhanced stimulus-evoked responsiveness. LTMRs exhibiting SF+ generally displayed increased firing during natural mechanical stimulation, consistent with spontaneous and evoked hyperexcitability arising from common mechanisms operating within sensory endings.

Although the molecular basis of this hyperexcitability remains unresolved, converging evidence provides reason to suspect that NaV1.6 channels contribute to these early abnormalities. OX prolongs NaV1.6-mediated inward currents in large-diameter sensory afferents (Siau et al., 2006). In previous studies, we demonstrated dense clustering of NaV1.6 channels at the heminode immediately adjacent to the sensory ending, the normal site of spike initiation in group Ia afferents [11] and subsequently showed through computational modeling that NaV1.6 exerts strong control over spindle excitability (Housley et al., 2024). Together with the present localization of spontaneous firing to distal axons adjacent to sensory endings, these observations raise the possibility that OX-induced enhancement of NaV1.6 activity provides a common mechanism linking spontaneous firing and enhanced mechanosensory responsiveness. Although NaV1.6 channels are also present at nodes of Ranvier (Duflocq et al., 2008), the distal localization of spontaneous firing argues that terminal rather than nodal channels contribute to the earliest abnormalities observed here.

Additional mechanisms are also likely to contribute. Sensory endings contain dense mitochondrial networks that support the substantial metabolic demands of continuous mechanotransduction (Zelená, 1994), and OX-induced mitochondrial dysfunction could impair ion homeostasis, depolarize terminal membranes, and increase the probability of spontaneous firing (Flatters et al., 2017; Illias et al., 2022; Xiao and Bennett, 2008; Xiao et al., 2012; Xiao et al., 2011). Other ion channel classes are also likely to contribute. Nevertheless, the convergence of previous evidence identifying NaV1.6 as a molecular target of OX, its strategic localization at sensory endings, its strong influence on sensory excitability, and the present localization of spontaneous firing to these same regions suggests that NaV1.6-mediated hyperexcitability represents an especially compelling mechanism for future investigation.

### Early sensory-ending dysfunction may precede later DRG pathology

The localization of spontaneous firing to sensory endings during the first day after OX exposure does not preclude later contributions from other excitable regions of the afferent. Voltage-gated sodium channels are concentrated not only at sensory endings but also within the axon initial segment of myelinated sensory neuron somata in the DRG. Computational modeling indicates that modest reductions in axon initial segment threshold can generate spontaneous firing (Nascimento et al., 2022), and numerous studies have implicated the DRG as a source of aberrant activity during chronic neuropathic states (Han et al., 2023; Lesniak et al., 2014; Michaelis et al., 2000; Zajaczkowska et al., 2019; Zhao et al., 2012). In contrast, our findings indicate that the DRG is not the primary source of spontaneous firing during the earliest phase of OX exposure.

This temporal sequence is consistent with recent observations in a mouse model of spared nerve injury, in which spontaneous firing emerged first at sensory endings and only later in DRG neurons (Zheng et al., 2022). Spontaneous activity arising from these two locations also appears to have different functional consequences. DRG-generated activity shows a stronger relationship to pain-related behaviors, and suppressing spontaneous cluster firing within the DRG abolishes pain behaviors more effectively than suppressing spontaneous activity in the peripheral nerve. These observations raise the possibility that sensory endings represent the initial site of dysfunction, whereas subsequent recruitment of the DRG marks an important transition toward clinically significant neuropathic symptoms.

Our findings capture events occurring within hours of OX administration, when structural abnormalities are already evident at sensory endings but before the broader transcriptional and network-level adaptations characteristic of established neuropathy. Consistent with this early time course, Jamieson and colleagues demonstrated rapid nucleolar pathology within DRG neurons after a single OX treatment, indicating that neuronal injury is already underway during the first day after exposure, although the source of spontaneous firing was not examined. Spontaneous firing typically emerged within 3–4 h of OX exposure in the present study, consistent with direct drug actions and rapidly developing metabolic consequences [32; 59; (Jamieson et al., 2005). Together, these findings suggest that spontaneous firing first emerges in the distal periphery, before evidence of recruitment of the DRG as a generator of aberrant activity. Although spontaneous firing initially originated at sensory endings, additional sites of spike initiation may emerge with time, as observed following peripheral nerve injury [74]. A similar progression during CIPN could ultimately include synchronized spontaneous activity among DRG neurons sufficient to engage central sensory circuits [12].

Consistent with this possibility, previous studies indicate that spontaneous firing eventually dissipates and is replaced by reduced mechanosensory responsiveness during the chronic phase of CIPN (Bullinger et al., 2011; Housley et al., 2021; Housley et al., 2022; Vincent et al., 2016). This transition from hyperexcitability to hypoexcitability raises the possibility of maladaptive homeostatic compensation and may help explain why the severity of acute OX-induced symptoms predicts the subsequent development and persistence of chronic neuropathy (Brownstone and Lancelin, 2018; Nascimento et al., 2024).

### Immediate consequences of spontaneous firing for sensory encoding and behavior

OX-induced spontaneous firing occurred at firing rates and temporal patterns sufficient to distort, mask, or generate false mechanosensory signals in individual LTMRs. These effects parallel observations in OX-treated mice and in patients with peripheral neuropathy (Campero et al., 1998; Saleque et al., 2024). In humans, brief bursts of activity in single tactile afferents evoke tingling or prickling sensations following transient ischemia or direct electrical stimulation (Macefield et al., 1990; Ochoa and Torebjork, 1980; Watkins et al., 2022). These observations provide a plausible mechanistic explanation for acute OX-induced paresthesias, even when spontaneous firing is present in only a minority of tactile afferents. Although subjective sensation cannot be assessed directly in rodents, similar activity in human LTMRs would be expected to generate tingling or prickling sensations and could potentially be detected using microneurographic recordings during acute OX treatment.

At the same time, the functional consequences of spontaneous firing appear surprisingly limited. Despite substantial disruption of spike timing in individual afferents, rats maintained normal performance of the hindlimb replacement strategy, an LTMR-dependent sensorimotor behavior. Likewise, population-level encoding remained largely preserved despite robust abnormalities in individual afferents. The predominance of SF− afferents together with the intermittent and asynchronous expression of spontaneous firing probably limited the impact of aberrant activity on ensemble signaling.

This dissociation helps explain how acute paresthesias can coexist with largely preserved function. Activation of individual tactile afferents is sufficient to evoke percepts, suggesting that spontaneous firing in these neurons may directly engage central sensory pathways underlying tingling or prickling sensations. In contrast, proprioceptive afferents and other LTMR subclasses, despite exhibiting spontaneous firing at a higher incidence, may fall short of disrupting sensorimotor function because too few afferents are affected and their activity remains poorly coordinated. More broadly, our findings indicate that abnormal activity distributed across tactile and proprioceptive LTMR submodalities is insufficient to disrupt all LTMR-dependent pathways. Instead, the functional consequences of spontaneous firing depend on how aberrant activity is integrated within specific central circuits, allowing some sensory pathways to be engaged while others remain largely unaffected. These findings demonstrate that even widespread spontaneous firing across tactile and proprioceptive LTMRs is not, by itself, sufficient to disrupt all LTMR-dependent sensory functions.

## Conclusions

Sensory endings are the earliest detected source of oxaliplatin-induced spontaneous firing and hyperexcitability across tactile and proprioceptive LTMR subtypes. Although spontaneous firing distorted spike encoding in individual afferents and may contribute to acute paresthetic symptoms, its limited incidence, intermittency, and asynchronous expression were insufficient to substantially disrupt population-level sensory coding or LTMR-dependent sensorimotor function. These findings identify sensory endings as an early target of oxaliplatin neurotoxicity, support a common peripheral mechanism for spontaneous and evoked hyperexcitability, and suggest that subsequent recruitment of the dorsal root ganglion may represent a critical transition in the development of chronic chemotherapy-induced peripheral neuropathy.

## Acknowledgements

This study was funded by National Institutes of Health R01CA268125 (TCC) and R01HD090642 (LHT).

All data are available on reasonable request to the corresponding author.

